# Characterization of microbial dark matter at scale with MetaSBT and taxonomy-aware Sequence Bloom Trees

**DOI:** 10.1101/2025.08.25.672238

**Authors:** Fabio Cumbo, Daniel Blankenberg

## Abstract

Metagenomics has become a powerful tool for studying microbial communities, allowing researchers to investigate microbial diversity within complex environmental samples. Recent advances in sequencing technology have enabled the recovery of near-complete microbial genomes directly from metagenomic samples, also known as metagenome-assembled genomes (MAGs). However, accurately characterizing these genomes remains a significant challenge due to the presence of sequencing errors, incomplete assembly, and contamination.

Here we present MetaSBT, a new tool for organizing, indexing, and characterizing microbial reference genomes and MAGs. It is able to identify clusters of genomes at all seven taxonomic levels, from the kingdom all the way down to the species level, using the Sequence Bloom Tree (SBT) data structure that relies on Bloom Filters (BFs) to index massive amounts of genomes based on their k-mers composition. We have built an initial set of databases composed of over 190 thousand viral genomes from NCBI GenBank and public sources grouped into sequence consistent clusters at different taxonomic levels, making it the first software solution for the classification of viruses at different ranks, including still unknown ones. This results in the definition of over 40 thousand species clusters where ∼80% do not match with any known viral species in reference databases to date. Furthermore, we show how our databases can be used as a new basis for existing quantitative metagenomic profilers to unlock the detection of unknown microbes and the estimation of their abundance in metagenomic samples. Finally, the framework is released open-source and, along with its public databases, is fully integrated into the Galaxy Platform enabling broad accessibility.

**Importance:** The MetaSBT framework and its databases, together with its integration in the Galaxy Platform, provide a powerful resource for microbial research. MetaSBT provides a powerful and scalable approach for classifying microbial genomes, including previously unknown ones. This facilitates the discovery and characterization of novel taxa, a crucial feature for expanding our knowledge of microbial diversity and its implications within host health and environmental factors. Furthermore, MetaSBT databases can serve as a reference base for other state-of-the-art tools, enhancing their capabilities to identify, analyze, and classify unknown microbes in metagenomic samples.

## INTRODUCTION

Microbial communities play essential roles in all aspects of life. They are involved in nutrient cycling, biogeochemical processes, and the maintenance of human health. However, our understanding of microbial communities is still limited. One of the main challenges is the identification and characterization of the vast majority of cultivation recalcitrant microbes.

Microbial genomics and metagenomics [1], the study of genetic material recovered directly from environmental samples, have revolutionized our understanding of microbial diversity and function, enabling us to identify and characterize new microbial species and communities without the need for cultivation [2], with profound implications for a wide variety of scientific fields, including medicine, agriculture, environmental science, and biotechnology. However, harnessing the full potential of large metagenomic datasets presents significant computational and analytical challenges. Accurately identifying and characterizing the myriad of microbes within complex environmental samples requires robust bioinformatics tools capable of handling massive datasets, resolving complex relationships between species, and accounting for the vast unknown diversity that still eludes our current understanding. One of the main challenges here is the clustering of genomes into taxonomically consistent groups.

Traditionally, microbial genomes have been clustered through methods that take into account their phylogeny, by inferring the evolutionary relationships between organisms and defining huge phylogenetic trees [3]. These are built by aligning genomes of different organisms and identifying similarities among their genomic regions. However, phylogenetic methods are not well-suited for clustering large datasets of genomes, mainly because they require the alignment of all the genomes under study, which can be computationally prohibitive, especially for datasets counting hundreds of thousands or even millions of genomes.

In recent years, a number of new methods have been developed for the clustering of microbial genomes, typically based on the analysis of k-mers, i.e., by comparing the set of k-mers present in each genome in the dataset. Genomes that share a large number of k-mers are supposed to be more closely related than genomes that share a small number of k-mers. However, existing k-mer based methods, such as the widely used species-level genome bins (SGB) approach based on the average-linkage hierarchical clustering of microbial genomes [2], rely on fixed thresholds to delineate clusters at different taxonomic levels (5% of genetic distance to define species, 10% to define genera, and 30% to define families). This approach, while valuable, can limit the ability to capture finer-grained taxonomic distinctions, especially for less well-characterized microbial lineages where the boundaries between clusters of genomes may be less clear-cut, especially at higher taxonomic levels where no thresholds can be easily and uniquely defined. Additionally, the thresholds on the genomes’ genetic distances have been designed to work with bacteria and archaea, while the same thresholds cannot be directly applied to other microbial kingdoms.

However, despite these limitations, the SGB-based clustering approach brought relevant advantages that led to the discovery of novel microbial taxa. For instance, a new bacterial species named *Cibionibacter quicibialis* [2] was identified and subsequently isolated. Moreover, it has been instrumental in refining our understanding of existing microbial groups. In a study focused on the *Segatella copri* [4,5] complex, formerly known as *Prevotella copri*, the SGB approach enabled the division of this group into 13 distinct species within the same genus. These findings underscore the ability of the SGB-based method to reveal intricate relationships within microbial communities and to redefine our understanding of microbial taxonomy.

Given the aforementioned problems of adopting current approaches and the undisputed benefits that it demonstrated to bring in advancing research in metagenomics, we built a scalable framework able to organize massive amounts of microbial genomes based on the latest algorithmic and data structure technologies. This is our solution in response to the lack of a systematic procedure to organize microbial genomes from isolate sequencing as well as metagenome-assembled genomes (MAGs) into sequence-consistent clusters.

MetaSBT is a new scalable k-mer-based framework specifically designed for the incremental clustering of massive amounts of microbial reference genomes and MAGs, with the final aim of uncovering new groups of yet-to-be-named microbes at different taxonomic levels, from the kingdom down to the species level. The development of MetaSBT was motivated by the need for a powerful and versatile tool for the organization of microbial genomes from different kingdoms and has the potential to revolutionize the way we study microbial communities.

This work describes the development and application of MetaSBT, focusing on its ability to:

- construct comprehensive, searchable, and incrementally updatable databases of microbial genomes;
- accurately classify microbial MAGs and genomes from isolate sequencing and organize them into taxonomically consistent clusters without fixed thresholds;
- facilitate the discovery and characterization of new microbial taxa;
- enhance the capabilities of quantitative taxonomic profilers by integrating our databases.

The rest of this manuscript is focused on (i) providing a detailed description of the MetaSBT framework, including its underlying methodology and data structures, (ii) constructing and validating MetaSBT databases, (iii) applying the framework on a set of public available viral genomes for building the first series of public MetaSBT databases and showing its ability to reveal novel taxa, (iv) exploiting our database to extend and improve the ability of quantitative taxonomic profilers to detect still-unknown viral species, and (v) integrating our framework and its databases into the Galaxy platform for broader accessibility.

## MATERIALS AND METHODS

The MetaSBT framework has been entirely implemented in Python 3.8 leveraging HowDeSBT [6], a software for building Sequence Bloom Trees (SBTs) [6–8], and extending its functionalities. SBTs are tree-like data structures whose nodes are Bloom Filters (BF), probabilistic data structures used to represent sets, representing one of the most efficient ways to index massive amounts of data to date [9]. SBTs have been thought specifically for indexing experimental sequence data, and BFs are usually built over sets of k-mers. In our case, we adopted this technology to index genomes. However, although HowDeSBT is quite fast and memory efficient, it comes with a significant limitation. Its unique way of encoding and clustering data, based on the concept of determined/how split filters [6], does not allow it to incrementally update the tree with new data once the tree is built, leaving the only option to rebuild the whole tree from scratch by extending the initial set of data (i.e., genomes in our case). Providing the ability to update the tree without affecting the efficient way it clusters data is a must-have feature since rebuilding everything from scratch every time we need to include a new genome in the database is clearly not feasible, especially when dealing with thousands of genomes which require a substantial amount of computational resources to process.

In order to overcome this issue, we propose to take advantage of the natural taxonomic classification of reference genomes from isolate sequencing. These, in fact, are well known and fully classified, from the kingdom all the way down to the species level. We used this information to build a series of SBTs, one for each taxonomic entity at all seven taxonomic levels (i.e., kingdom, phylum, class, order, family, genus, and species) as described in the next section.

MetaSBT modules are briefly introduced in Box 1 and described in depth in the following sections, while the whole framework architecture is reported below in Figure 1.

**Figure 1:**
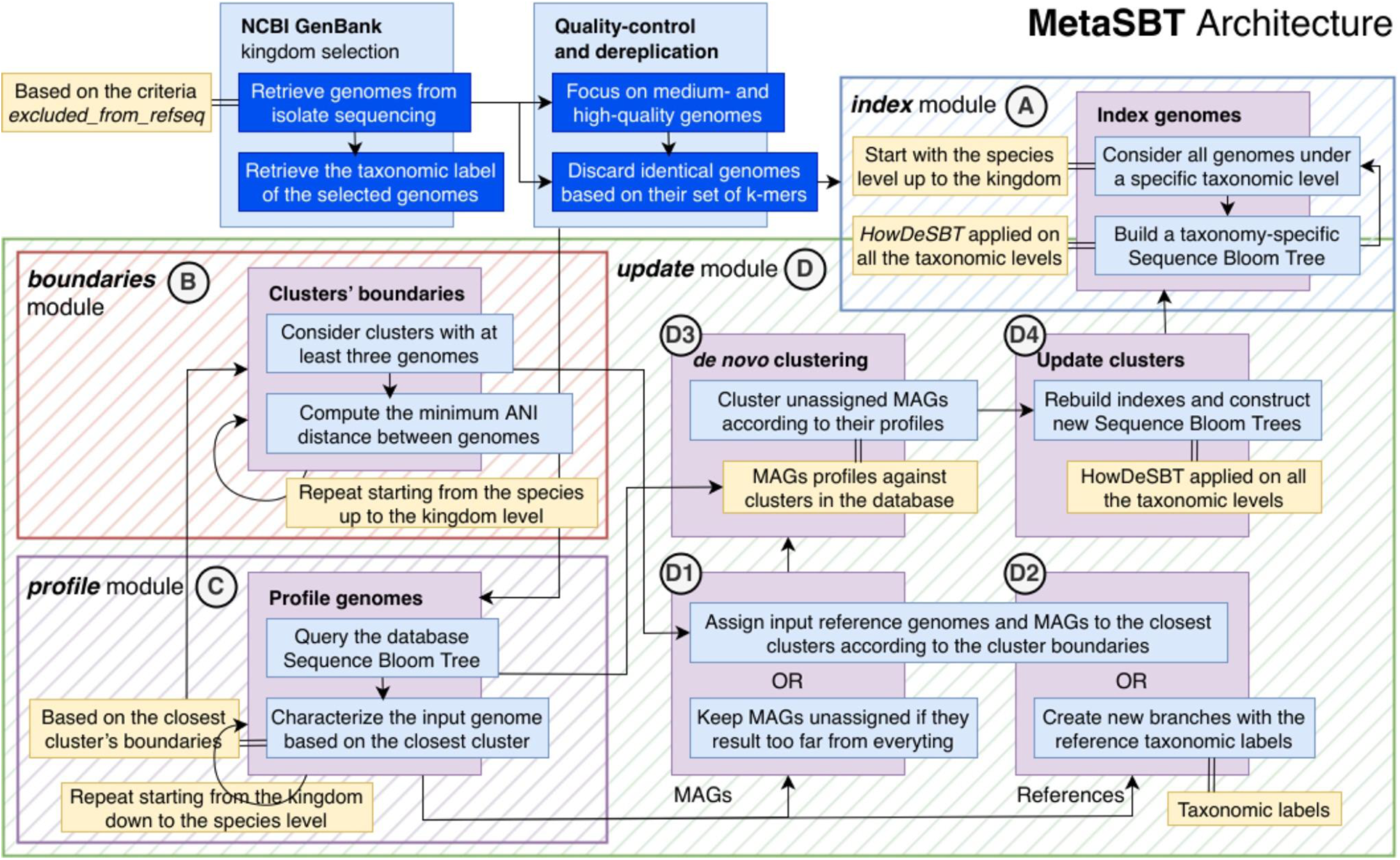
The MetaSBT framework architecture is mainly composed of 4 modules, i.e., *index*, *profile*, *boundaries*, and *update*. The *index* module (box **A**) works as a wrapper around HowDeSBT and it is used to build a series of SBTs for each of the clusters at every taxonomic level, starting from the species up to the kingdom; the *boundaries* module (box **B**) defines the clusters’ boundaries defined as the minimum and maximum ANI distances between all the genomes under the same cluster; the *profile* module (box **C**) is used to characterize a genome at all the seven taxonomic levels according to the closest clusters in the database tree and their boundaries; finally, the *update* module (box **D**) provides a method to incrementally update a SBT-based database by creating new clusters with new sets of MAGs and eventually characterize previously unknown clusters with new sets of reference genomes.

### Box 1: MetaSBT framework features as functional modules

- ***index*:** this is responsible for indexing a set of reference genomes, often retrieved from NCBI GenBank, by organizing them following their taxonomic classification, and building a series of SBTs at all taxonomic levels (see Figure 1 box A). It also support a quality control step and a dereplication procedure to make sure to exclude low-quality genomes and avoiding processing the same genomes twice;
- ***boundaries*:** this processes every cluster on all seven taxonomic levels and reports the clusters boundaries as the minimum and maximum Average Nucleotide Identity (ANI) distances computed among all the genomes under a specific cluster (see Figure 1 box B);
- ***profile*:** the framework also provides a profile module responsible for the characterization of input genomes at all the taxonomic levels by querying the database and reporting the distance between the input genome and the closest clusters (see Figure 1 box C);
- ***update*:** this is responsible for updating the database with new genomes. First, it makes use of the profile module to establish the closest kingdom, phylum, class, order, family, genus, species, and genome in the database from the input genomes (see Figure 1 box D, steps D1–4). Here is where the results of the boundaries module are required in order to establish, together with the result of the profile module, whether an input genome should be assigned and added to the closest cluster by looking at the ANI distance between the input genome and the closest clusters at all the seven taxonomic levels.

### Exploiting the natural taxonomic classification of reference genomes is the key to efficiently update a SBT-based database

Here we propose a way for decomposing the SBT data structure into a series of SBTs exploiting the taxonomic organization of reference genomes. This process starts by grouping genomes belonging to the same species, and producing a SBT over their set of k-mers. This produces a tree for each of the species involved in the initial set of genomes. The root nodes of the species trees are then used as leaves for building the SBTs at the genus level, by grouping species belonging to the same genus according to their taxonomic classification. Note that here we group the root nodes of the species SBTs that contain the whole genetic content of all the genomes under those specific trees. Thus, considering the root nodes is actually like considering the whole set of genomes used to build the species trees. The process continues iteratively, by considering the root nodes of the genera trees as leaves for building the SBTs at the family level, and analogously for the order, class, phylum, and finally the kingdom level composed of a single SBT only whose root node contains the genetic content of all the genomes in the initial set of genomes. Adopting this technique allows updating the tree as follows: given a new genomes and once the closest genome in the database has been established, we should first determine whether the closest cluster (i.e., the species of which the closest genome belongs to) is close enough to eventually add the input genome to it (see next section). Here we have two options:

- in case the input genome has resulted to be close enough to the closest species, we reconsider the whole set of genomes belonging to that species plus the input genome to rebuild the species cluster. Then we reconsider the root nodes of all the species clusters belonging to that specific genus to rebuild the genus tree, and so on, repeating the same steps used to initially build the database. The difference here is that <u>we are rebuilding a series of SBTs, one for each of the seven taxonomic levels, corresponding to a single branch of the whole SBT database</u>;
- otherwise, if we determine that the input genome is not close enough to the closest species in the database, we can then define a new species, as soon as the input genome results to be close enough to the closest genus tree, and so on. This ends up in defining new clusters at different taxonomic levels that, in case of updating the database with MAGs, do not contain any reference genomes at all, representing still-unknown species. This process is described in depth in the next sections.

### The size of the database is directly proportional to the length of the Bloom Filters

SBTs are tree-like data structures whose nodes are Bloom Filters (BFs), i.e., probabilistic data structures used to represent sets. In our specific case, BFs are used to represent the set of distinct k-mers in a particular genome. The numerosity of k-mers also determines the size of the BFs, usually represented as bit vectors where 1 denotes the presence of a specific k-mer, while 0 represents its absence. K-mers are indeed encoded into 1s under specific positions in the BF vectors that are established as the result of one or more hash functions applied on each k-mer.

Establishing a proper BF size is a key aspect in the definition of the whole SBT database. The larger the BFs, the more storage space is required to maintain these data structures and the whole database, but smaller BFs translate in high false positive rates affecting their lookup capabilities. Thus, in order to estimate a proper BF size and also minimize their memory footprint, we propose to estimate the optimal BF size for each taxonomic level, based on the total number of k-mers belonging to all the genomes under that specific level. Using a universal size for all the BFs in the database would indeed produce very sparse BFs, especially at the lower end of the taxonomic organization, potentially leading to a significant waste of storage space.

Instead, we estimate the BF size at the species level by summing up the number of distinct k-mers from all the genomes belonging to each species. At the genus level, we estimate the BF size by summing up the number of distinct k-mers from all the species belonging to each genus. This process is repeated iteratively up to the kingdom level where the whole set of genomes in the database is taken into consideration to establish the BF size.

This way, we define a different BF size for each taxonomic level, which takes into account the different number of k-mers that is stored at each level. Depending on the number of genomes initially considered to build the database, this technique allows drastically reducing the storage space required to maintain the whole database tree.

Also note that the number of distinct k-mers strictly depends on their length. A larger k-mer size might also lead to larger BFs, impacting the storage requirements, as we previously mentioned, in addition to the time required to query the data structures. Therefore, selecting an appropriate k-mer length is also crucial for balancing database size and performance. To address this, MetaSBT integrates *KITSUNE* [10], a Python tool which enables the automatic estimation of an empirically optimal k-mer size based on a combination of the Cumulative Relative Entropy, Average Number of Common Features, and Observed Common Features measures computed on the initial set of genomes across a specific range of k-mer sizes.

However, it is worth to note that the strategy mentioned above also comes with the side effect of a faster saturation of the BFs at the lower taxonomic levels. While deciding whether applying this procedure or not is a crucial first step in building a database, we can also mitigate the side effects, as described in the following paragraphs.

### Preventing a fast saturation of Bloom Filters: the larger the better

A critical aspect of MetaSBT’s performance and efficiency lies in the initial evaluation of a proper BF size computed as an estimation of the total number of unique k-mers in the union of all the initial set of genomes minus the k-mers that occur only once. Note that discarding single-occurrence k-mers is optional, but it is usually advised since they could be the result of sequencing errors. This estimation is performed with the support of the k-mer counting tool *ntCard* [11].

Since our goal is to build a database that can scale to accommodate an ever-growing number of newly sequenced genomes, we have to deal with the issue of the saturation of the BF: as more genomes are added to the database over time, the BF might become saturated, eventually affecting the overall system’s performance and database, up to the point that inserting new genomes is no longer possible, especially in the case of genomes whose content is different from the existing ones. More specifically, BF saturation occurs when a significant portion of the filter’s bits are set to 1. This increases the likelihood of false positives, where the filter might incorrectly indicate the presence of a k-mer that was never actually added. Thus, determining a proper BF size upfront is key to ensure the long-term scalability of MetaSBT without having to completely rebuild the database from scratch.

Please note that reaching the complete saturation of the BFs is the only extreme case that we must avoid by simply incrementing the *ntCard* estimation largely enough to account for future growth of the database.

### How close is close enough? Defining dynamic clusters’ boundaries

In order to profile a genome and eventually update a database, we should determine whether the input genome is close enough to its closest species cluster in the database to be characterized as that specific species. In order to do so, we introduce the concept of cluster boundaries defined as the minimum and maximum ANI distance between all the genomes under a particular cluster. These numbers are computed for all the clusters at all the seven taxonomic levels, and give us an idea of the overall genetic diversity between genomes in the database. At the time of deciding whether a new genome must be added to the closest cluster, we then compare the ANI computed between the genome and the closest cluster, and we finally compare it with the closest cluster’s boundaries. This ends up in eventually rebuilding the cluster or defining a new one as mentioned earlier.

Please note that, in order to define the cluster boundaries, the cluster must contain at least three genomes. Thus, if a cluster contains less than three genomes, we implicitly deduct its boundaries as the average minimum and maximum of all the other clusters under the same upper taxonomic level. This is, however, just an approximation and a temporary solution, and the boundaries of these clusters must be recomputed with future incremental updates, as soon as they will contain at least three genomes. At that point, all the MAGs that have been previously assigned to that cluster but now are out of its boundaries, must be reprocessed and reclustered, leading, eventually, to the generation of new clusters as discussed below.

### A tree structure to speed-up the characterization of genomes

Once we build a first database with reference genomes only and we computed the clusters’ boundaries, we are already able to profile new genomes, which is also a step required to update the database. Note that the first baseline must contain reference genomes only since we use their taxonomic classification to define clusters at all the seven taxonomic levels.

In order to profile a new genome we exploit the tree structure to initially query the root node of the kingdom tree with the set of k-mers in the input genome. The query reports the number of k-mers in common between the input genome and the kingdom. This process is repeated another six times by querying the root nodes of all the other clusters at the remaining taxonomic levels, up to the species level. Note that before proceeding to query a specific cluster and proceed to the next taxonomic level, we first prune the database tree by considering a subset of clusters selected by applying a distance threshold based on the number of k-mers matching with the closest cluster plus its 25%. Expanding the set of closest genomes by a 25% in genetic distance can result in querying way more branches of the database tree, and thus taking more time to establish the closest nodes, but it is also crucial for mitigating, and sometimes preventing, misclassifications due to the problem of horizontal gene transfer between distantly related species, e.g., in case of human-associated microorganisms [12].

Upon reaching the species level, we finally profile the input genome with the same taxonomy as the closest species, only in case the ANI between the input genome and the closest species falls within the closest species’ boundaries. This process also reports the closest genome under the selected closest species, which is also the closest strain.

### Beyond known taxonomies: MetaSBT’s approach to defining new microbial taxa

While established taxonomic classifications provide a valuable framework for understanding microbial diversity, they are inherently limited by our current knowledge. Metagenomic studies constantly reveal novel microbial lineages that defy easy categorization within existing taxonomic ranks. This section explores how MetaSBT goes beyond known taxonomies, employing a clustering-based approach to identify and define new microbial taxa.

To achieve this goal, MetaSBT needs to start from a baseline of known microbial reference genomes, i.e., genomes from isolate sequencing characterized at all the seven taxonomic levels, from the kingdom down to the species level. As we previously discussed, MetaSBT exploits their natural taxonomic classification to build a multitude of SBTs at all the seven known taxonomic levels, producing the first baseline of the MetaSBT database. Cluster boundaries are then computed in order to provide users with an initial overview of the intra-clusters genetic diversity. These are also crucial for establishing, in conjunction with the profiles (see previous sections), whether a new MAG should be assigned to the closest cluster or not.

Starting from the species level, in order to assign a MAG to its closest species according to its profiles, MetaSBT does not directly look at the ANI between the input genome and the closest genome in the database. That is because the closest genome could be closer to the boundaries of its own cluster compared to other genomes in the same species. Thus, MetaSBT proceeds by computing the ANI between the input genome and the centroid of the closest species (which could eventually overlap with the closest genome), defined as the genome that maximizes the ANI similarity with all the other genomes under the same cluster. This is finally compared with the species boundaries to eventually assign the genome to that particular species.

This process runs iteratively on all the input genomes, by keeping apart those that do not fall within the boundaries of any species clusters in the database. If the process ends up in assigning a genome to its closest species, it then immediately performs an average-linkage hierarchical clustering on the pair-wise ANI matrix computed on all the input genomes to finally use the boundary of the closest species as threshold for the identification of other genomes to be assigned to the same species.

The same procedure is applied downwards from the species level up to the kingdom level, using different thresholds based on the actual boundaries of the input genomes’ closest clusters. Note that no new clusters have been created yet, and all the input genomes have been assigned to a lineage with different levels of knowness.

In order to complete the definition of the assigned taxonomies, MetaSBT starts processing the input genomes by grouping them according to their partial common labels, from the kingdom upwards to the species level. Since these genomes did not fall within the boundaries of any clusters under this specific taxonomic level, MetaSBT estimates the boundaries of the new clusters as the average between the boundaries of all the other clusters under the same taxonomic level. The estimated boundary is used as threshold for the definition of new clusters upon the computation of the average-linkage hierarchical clustering on the partially characterized genomes with the same common partial taxonomic label. The same process is repeated multiple times until a complete characterization of all the input genomes with at least one level of unknownness.

This process ends up in defining new clusters at different taxonomic levels which, by definition, do not contain any reference genomes at all, thus representing still unknown microbial characterizations.

It is worth noting that, in case of new reference genomes, we do not rely solely on their already known taxonomy. We instead apply the same process performed on MAGs but, at the time of assigning new references to already defined clusters in the database, their taxonomies contribute to the definition of the taxonomic characterization of the assigned clusters. In case the assigned cluster is unknown, it inherits the taxonomic profile of the new references, thus losing its unknown traits. Otherwise, the taxonomic labels of the new reference genomes, together with the taxonomies of all the other references in the same assigned cluster, contribute to the majority voting mechanism used to pick the most occurring taxonomy to characterize the cluster.

### Database integrity: quality control and dereplication

As the database grows with new genomes, we need to ensure its integrity by performing quality control and dereplication on the incoming genomes before adding them to the database.

Quality control of input genomes is a necessary step to validate the completeness and contamination of genomes. MetaSBT integrates different quality control frameworks to assess genome quality depending on their kingdom. More specifically, it integrates *CheckV* [13] for viral genomes, *CheckM* [14] for bacterial and archaeal genomes, and *BUSCO* [15] for eukaryotic genomes.

MetaSBT also provides a dereplication procedure to ensure no duplicate genomes are processed and included twice in the database. We set a dereplication threshold at 99% of similarity between genomes by default based on their ANI distance.

### Standardizing genomes and samples metadata: a powerful tool in support of large-scale comparative genomic studies

While genomic sequences provide a fundamental understanding of microbial communities, their full interpretative potential is realized when considered in conjunction with their metadata, including information about their samples.

More specifically, sample metadata encompasses the contextual information surrounding sample collection, offering crucial insights into the ecological and experimental factors shaping microbial diversity and function.

Such metadata can encompass a wide range of details, including:

- *collection date and geographical location*: providing spatiotemporal context to understand species distribution and potential environmental influences;
- *environmental conditions*: factors like temperature, pH, salinity, nutrient availability, and more, offering insights into the ecological niches of different species;
- *host information*: for microbes associated with plants, animals, or other organisms, this data can reveal symbiotic relationships, pathogenicity, and host-specific adaptations;
- *sequencing technology and parameters*: ensuring transparency and reproducibility by documenting the methods used to generate the genomic data;
- *experimental conditions*: controlled experimental variables, such as treatment groups, sampling time points, and sequencing technologies employed, are crucial for interpreting observed patterns in microbial communities.

A pivotal component of this framework is the MetaSBT Data Model, which has been designed to ensure robust organization and standardization of genomes and samples metadata. Our data model is indeed composed of two main core components defined as JSON schemes [16]:

- the *genome* model component has been designed to keep track of genomes, sequences, and MAGs information like:

○ *metasbt_id*: a genome’s unique identifier assigned internally by MetaSBT;
○ *genome_id*: the original name or identifier assigned to the input genome before being processed by MetaSBT;
○ *sample_id*: the name or identifier of the metagenomic samples where a genome comes from;
○ *dataset_id*: the name or identifier of a dataset defined as a collection of one or more metagenomic samples;
○ *genome_type*: the nature of an input genome (i.e., reference genome or MAG);
○ *taxonomy*: the taxonomy assigned by MetaSBT to the cluster that a genome belongs to, that could potentially present some level of unknownness;
○ *completeness*, *contamination*, and *strain_heterogeneity*: a few information about the quality of the input genome.
- the *sample* model component, design to collect and standardize information like:

○ *sample_id*: the name or identifier of the metagenomic sample where reference genomes and MAGs come from;
○ *dataset_id*: the name or identifier of a dataset defined as a collection of one or more metagenomic samples;
○ *ecosystem*, *ecosystem_category*, *ecosystem_type*, *ecosystem_subtype*, and *specific_ecosystem*: attributes to categorize samples according to the environment where they have been collected from;
○ *sequencing_platform*: the platform used to sequence the metagenomic sample;
○ *age* and *age_category*: the age of the subject where the metagenomic sample has been collected from;
○ *gender*: the biological gender of the donor;
○ *country*: the country of origin of the donor;
○ *location*: the specific geographical location where the donor used to live at the time of collecting the metagenomic sample;
○ *disease*: whether the donor is affected by a specific disease at the time of collecting the metagenomic sample.

Note that we listed and discussed the most relevant attributes of the MetaSBT Data Model. Also note that our selection of metadata partially integrates the standardized attributes of genomes and samples metadata in the *curatedMetagenomicData* package [17] for R and those used in *JGI GOLD* [18] (Genomes Online Database of the Joint Genome Institute, Lawrence Berkeley National Laboratory). Please refer to the official MetaSBT documentation for additional information about the complete set of standardized attributes (see Availability section below).

Integrating sample metadata with genomic data enables researchers to move beyond taxonomic classification and delve into the ecological and evolutionary dynamics driving microbial community structure and function.

Keeping track of genomes and samples metadata has the potential to transform MetaSBT into a powerful tool for large-scale comparative genomic studies. Researchers can leverage this information to move beyond simple taxonomic classifications and delve into the intricate interplay between microbial communities and their surrounding environments.

For example, imagine discovering a cluster of previously unknown viral sequences within a dataset. By analyzing the associated metadata, researchers might find that these viruses consistently appear in samples collected from a specific geographical region or within a particular host species. This could point to unique ecological interactions or even potential zoonotic risks.

Furthermore, correlating metadata with genomic variations within a species could uncover adaptive traits linked to specific environmental pressures. This has profound implications for understanding how microbial communities evolve and respond to changing conditions, a crucial area of study in the face of climate change and other global challenges.

## RESULTS

We built the first baseline of the database considering all the viral reference genomes in NCBI GenBank [19]. The set of references comprises 26,285 genomes organized into 17 phyla, 38 classes, 63 orders, 161 families, 1,014 genera, and 8,169 species. Genomes are automatically retrieved with the *get_ncbi_genomes* subroutine implemented in MetaSBT that establishes whether a genome in NCBI GenBank is a reference genome or a MAG according to a series of rules. We focused on the attributes under the *excluded_from_refseq* metadata in the NCBI Assembly Report to discriminate reference genomes and MAGs.

In particular, a genome is defined as reference if its attributes are all under the list in Table 1 below.

**Table 1:**
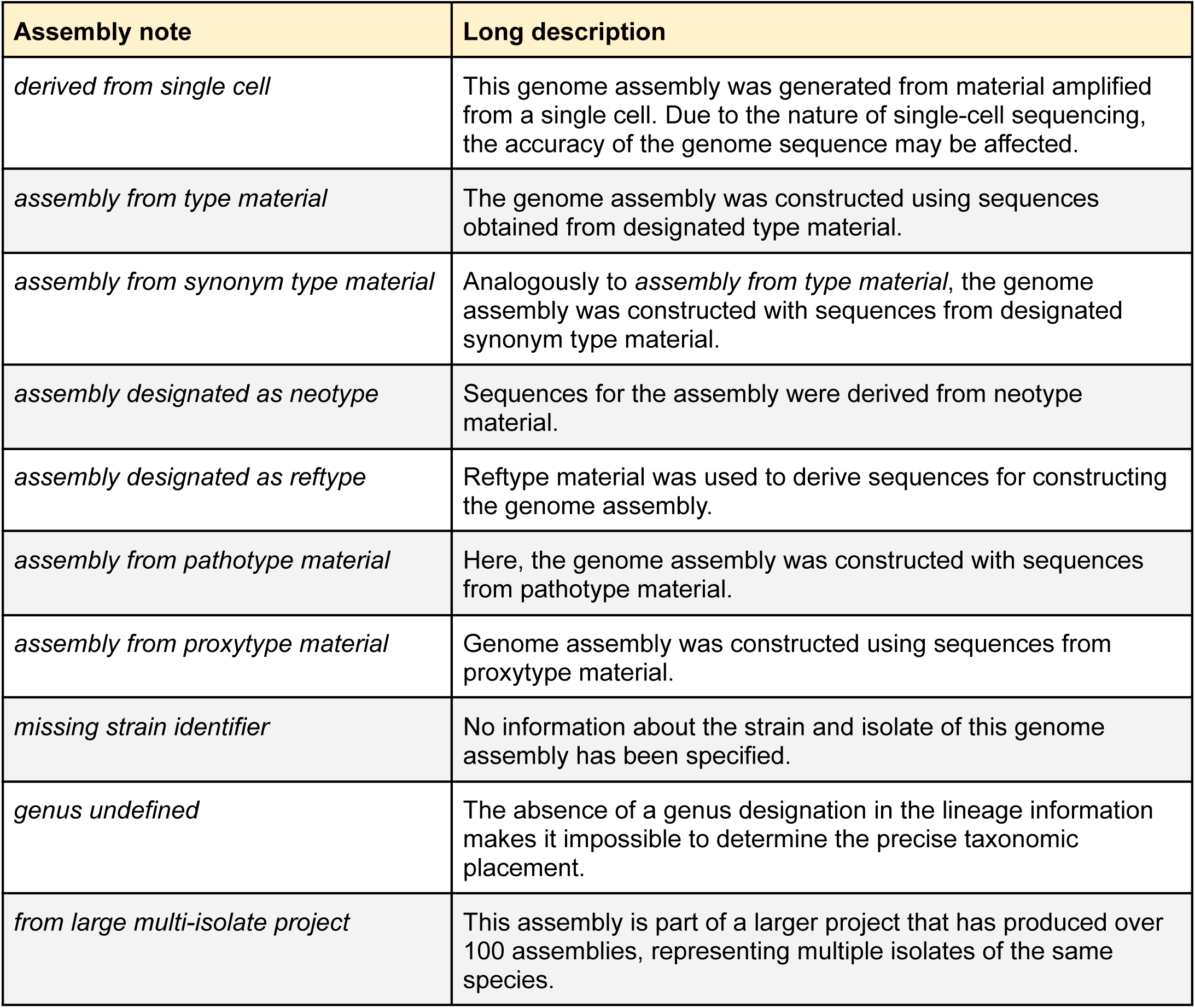
NCBI RefSeq exclusion criteria used to identify assemblies in GenBank as reference genomes in MetaSBT.

| Assembly note | Long description |
| --- | --- |
| *derived from single cell* | This genome assembly was generated from material amplified from a single cell. Due to the nature of single-cell sequencing, the accuracy of the genome sequence may be affected. |
| *assembly from type material* | The genome assembly was constructed using sequences obtained from designated type material. |
| *assembly from synonym type material* | Analogously to *assembly from type material*, the genome assembly was constructed with sequences from designated synonym type material. |
| *assembly designated as neotype* | Sequences for the assembly were derived from neotype material. |
| *assembly designated as reftype* | Reftype material was used to derive sequences for constructing the genome assembly. |
| *assembly from pathotype material* | Here, the genome assembly was constructed with sequences from pathotype material. |
| *assembly from proxytype material* | Genome assembly was constructed using sequences from proxytype material. |
| *missing strain identifier* | No information about the strain and isolate of this genome assembly has been specified. |
| *genus undefined* | The absence of a genus designation in the lineage information makes it impossible to determine the precise taxonomic placement. |
| *from large multi-isolate project* | This assembly is part of a larger project that has produced over 100 assemblies, representing multiple isolates of the same species. |

In case one or more attributes are not in the list among those in Table 1, we check whether the genome must be discarded according to the list of attributes in Table 2 below.

**Table 2:** NCBI RefSeq exclusion criteria used to discard assemblies from GenBank in MetaSBT.

| Assembly note | Long description |
| --- | --- |
| *abnormal gene to sequence ratio* | The number of genes predicted by NCBI PGAP for this assembly falls outside the typical range, suggesting potential overprediction or underprediction. The gene-to-sequence ratio, computed as the number of genes per kilobase of sequence, typically falls between 0.8 and 1.2. In this case, the ratio deviates from the expected range of 0.5 to 1.5, indicating an atypical gene density. |
| *chimeric* | The genome assembly appears to be contaminated, containing sequences from two distinct organisms that have been assembled together. |
| *contaminated* | The genome assembly contains extraneous sequences, potentially including material from other organisms, cloning vectors, linkers, adapters, or primers. |
| *genome length too large* | The assembled genome size significantly exceeds the expected size range for a given species, potentially due to contamination or assembly errors. |
| *genome length too small* | Analogously to *genome length too large*, the genome size is significantly smaller than the expected size range for a given species, raising concerns about the completeness of the assembly. |
| *hybrid* | The genome assembly appears to be derived from a hybrid organism, combining genetic material from different species, strains, or isolates. |
| *low gene count* | The NCBI PGAP gene prediction for this assembly yielded a significantly lower gene count than expected based on comparisons with high-quality reference genomes from the same species. |
| *low quality sequence* | Significant portions of the genome assembly exhibit characteristics of low sequence quality, such as long stretches of ambiguous bases, low-complexity regions, or other indicators of potential sequencing or assembly errors. |
| *many frameshifted proteins* | The genome assembly has an unusual high proportion of protein-coding genes with frameshift mutations, exceeding acceptable thresholds based on either a species-specific average (three standard deviations or 5% of annotated genes, whichever is greater) or a general threshold of 30% for species with limited genomic data. This high frameshift rate suggests potential sequencing errors, assembly errors, or biological factors impacting the organisms' gene expression. |
| *metagenome* | The assembly appears to originate from a metagenomic sample, containing sequences from a mixture of unidentified organisms, rather than a single, isolated species. |
| *misassembled* | Alignment to related genome assemblies or other available evidence suggests the presence of errors within this genome assembly. |
| *mixed culture* | This genome assembly is derived from a sample containing a co-culture of multiple organisms, rather than a single, isolated |
|  | species. |
| *untrustworthy as type* | This refers to situations where the genome assembly is deemed unreliable or questionable for a multitude of reasons. |

If a genome is not recognised as a reference and it is not discarded according to the list of attributes in Tables 1 and 2 respectively, it is considered a MAG (please refer to the official NCBI documentation for additional details about the assembly metadata used to discriminate genomes in MetaSBT at https://www.ncbi.nlm.nih.gov/assembly/help/anomnotrefseq/).

We also retrieved the set of 1,111 MAGs from NCBI GenBank, where genomes are classified as MAGs according to the same set of rules reported above. We used these genomes to produce a new database as the result of updating the initial database with reference genomes only. The update process ended up with the definition of 28 new orders, 79 new families, 268 new genera, and 936 new species clusters.

In order to demonstrate the MetaSBT ability to handle heterogeneous sources, we also considered the viral sequences distributed in the public Metagenomic Gut Virus catalogue [20] (MGV) over the MetaSBT database with NCBI GenBank genomes and MAGs as an incremental update. Note that MGV contains a metagenomic compendium of roughly 190-thousand DNA viruses from the human gut microbiome that we treated as MAGs for updating our database. It greatly expanded the set of unknown taxa to 2,976 new classes, 6,460 new orders, 8,622 new families, 12,634 new genera, and 31,624 new species. Note that the MGV genomes were originally identified using a comprehensive viral detection pipeline developed by the MGV team, which integrates multiple features and classifiers to distinguish viral contigs from non-viral ones in metagenomic assemblies. The pipeline combines gene prediction via Prodigal [21], functional annotation through HMMER [22] searches against both viral (IMG/VR) and microbial (Pfam) marker gene databases [23,24], k-mer based viral sequence scoring using VirFinder [25], and sequence-level characteristics like strand switch rates. The final classification step is based on a rule-based system that integrates these features using logic that is adapted to different contig length ranges. This pipeline is well explained in the MGV publication [20] and in GitHub at https://github.com/snayfach/MGV. We are currently planning to adopt the same procedure in future analyses of publicly available metagenomic samples. The viral genomes recovered from these them will be periodically integrated into MetaSBT.

Also note that the MetaSBT database presented here has been constructed considering the whole set of k-mers in the involved genomes. As previously explained, we used *KITSUNE* that reported 9 as the best k-mer size for our set of genomes. This parameter is also used to establish a proper BF size, which is determined with *ntcard* as the total number of distinct k-mers, aso considering k-mers with a single occurrence in this very specific context (i.e., Viruses). Note that we increase the estimated BF size by 100% in order to make the BFs big enough for future updates and avoid early saturation problems as previously discussed.

Since the k-mer size is involved in every aspect of the MetaSBT framework, once established it can only be changed at the price of rebuilding the whole database from scratch. This is why it is so important to estimate a proper kingdom-specific k-mer size before building a database.

Finally, Figure 2 shows the distribution of 40,729 MetaSBT clusters at the species level in descending order based on their number of genomes, comprising 8,169 known (∼20% – red) and 32,560 unknown (∼80% – black) species. We limit the number of clusters in the barplot to species with >100 genomes (39 known and 207 unknown clusters), thus focusing on the 0.6% of the clusters in out database, while we report a histogram in the upper-right portion of the figure with the number of known and unknown clusters considering only the species with a number of genomes ≤100 (8,130 known and 32,353 unknown clusters), with a number of single-genome clusters equals to 25,579 (7,192 known and 18,387 unknown clusters). Please look below for a discussion about the reliability of new clusters.

**Figure 2:**
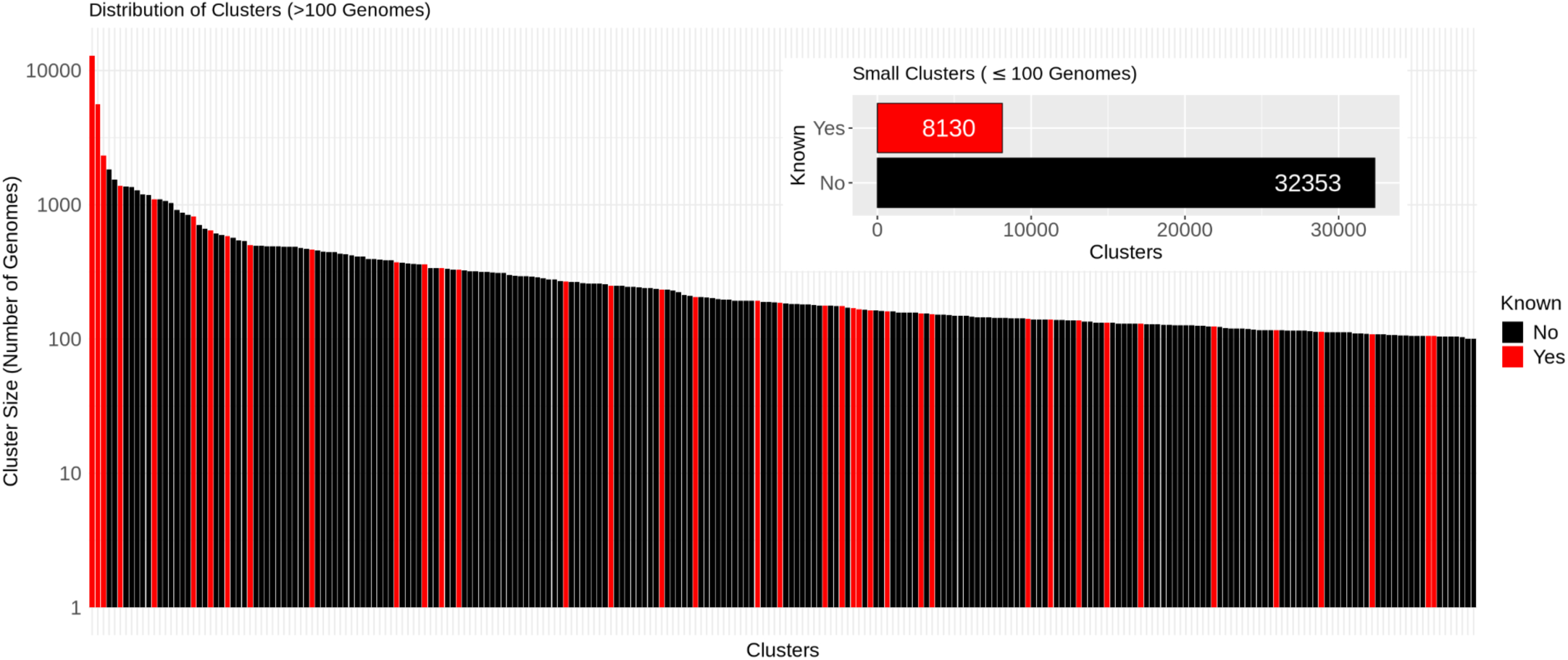
Color-coded distribution of 8,169 known (red) and 32,560 unknown (black) clusters at the species level comprising 26,285 reference genomes and 190,756 MAGs from the NCBI GenBank and MGV repositories. Clusters with a number of genomes >100 (39 known and 207 unknown) are sorted in a descending order based on their size, while the histogram reports the number of known and unknown clusters with ≤100 genomes, i.e., 8,130 and 32,353 respectively.

Analogously, considering a total number of 13,916 genera, 1,014 are known (∼7.3%), while 12,902 are unknown (∼92.7%). At the family level, considering a total number of 8,862 clusters, 161 are known (∼1.9%), while 8,701 are unknown (∼98.1%). The order level is instead composed of 6,551 clusters, of which 63 are known (∼1%), while 6,488 are unknown (∼99%). With 3,014 classes, only 38 clusters are known (∼1.3%), while 2,976 are still unknown (∼98.7%). Finally, all of the 17 phyla defined in MetaSBT are known.

A summary of the MetaSBT Viruses databases as incremental updates is also reported in Table 3 below, while the list of known and unknown clusters at all the seven taxonomic levels, with their number of reference genomes and MAGs, BF density, and mean completeness and contamination, is provided in Supplementary Spreadsheet S1.

**Table 3:** Incremental updates of the public MetaSBT database with viral reference genomes and MAGs from NCBI GenBank and the MGV repository. For each of the databases, we show their data source, the number of genomes, in addition to the number of known and unknown clusters at the species level produced by MetaSBT. Note that the number of genomes is incremental, thus, if the number of reference genomes remains constant compared to the previous update, no new reference genomes have been processed (e.g., Reference Genomes in *20250115* → *20250118*). For the same reason, the actual number of new MAGs considered in an update must be computed as the number reported at that update minus the number of MAGs in the previous update (e.g., the actual number of MAGs processed in *20250320* is 189,645).

| Database ID | Data Source | References | MAGs | Known sp. | Unknown sp. |
| --- | --- | --- | --- | --- | --- |
| 20250115 | NCBI GenBank | 26,285 | 0 | 8,169 | 0 |
| 20250118 | NCBI GenBank | 26,285 | 1,111 | 8,169 | 936 |
| 20250320 | MGV | 26,285 | 188,240 | 8,169 | 32,560 |

Please note that, although MetaSBT integrates quality control and dereplication techniques, we did not perform a check on the quality of genomes and sequences in our database on purpose at this stage, neither a dereplication, for the following reasons:

1. we did not dereplicate genomes, especially MAGs, to allow unknown clusters to grow over time with subsequent incremental update and potentially confirm those growing clusters as actual novel taxa. This allows eventually discriminating actual taxa from artifacts (see dedicated section below);
2. we considered all the reference genomes and MAGs regardless of their quality in order to eventually allow users to identify junk sequences among their set of input sequences.

Although we did not perform any quality control procedures so far, we provide the complete list of viral genomes and sequences considered in our study, together with their quality in terms of completeness and contamination computed by *CheckV*, in Supplementary Spreadsheet S2.

Also note that MetaSBT databases are fully reproducible and the pipeline used to generate them in an incremental way is available and documented on GitHub at the MetaSBT-DBs repository (see section Availability for additional details).

### Misclassification of genomes in NCBI GenBank: the need for a new approach

The National Center for Biotechnology Information’s GenBank database stands as a cornerstone of genomic research, providing a vast repository of genetic information. However, despite its invaluable contributions, GenBank faces an ongoing challenge: the misclassification of genomes. This issue arises from a variety of factors, including the inherent complexities of biological classification, evolving taxonomic standards, and the constantly increasing volume of data deposited everyday. Misclassifications can have cascading effects, propagating errors in downstream analyses, hindering comparative genomics studies, and potentially leading to inaccurate conclusions. For instance, a misclassified viral genome within a database could lead to the misidentification of related viruses in new metagenomic samples, obscuring true biodiversity patterns and hindering efforts to understand viral evolution and ecology. This underscores the urgent need for new approaches to genome classification that are robust, scalable, and adaptable to the evolving landscape of genomic data. MetaSBT, with its emphasis on comprehensive genomic comparison and its ability to incorporate updated taxonomic information, offers a promising way to address these challenges and improve the accuracy and reliability of genome classification.

Since the taxonomic information of reference genomes in NCBI GenBank is used to define the baseline of the MetaSBT databases, we implemented a clustering-based approach to refine the taxonomic assignments of viral reference genomes, using our definition of reference as previously discussed.

Our methodology involves the following steps:

1. <u>clustering of reference genomes</u>: we performed an average-linkage hierarchical clustering of all viral reference genomes retrieved from NCBI GenBank. Since we are dealing with Bloom Filters as sketch representations of genomes, we used the Jaccard index as a measure of genetic similarity. Considering two genomes A and B, the Jaccard index is defined as 1 minus the cardinality of the intersection over the cardinality of the union of A and B’s sets of unique kmers, as reported in Formula 1 below.

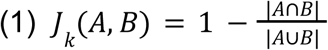 Then, we estimate the ANI between A and B without the need for an exact alignment, by applying the formula reported below (see Formula 2 where *k* is the k-mer size and *J_k_*is the Jaccard index as reported in Formula 1) [26–29]:

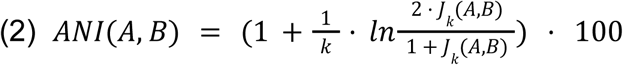
2. <u>cluster boundary definition</u>: clusters were delineated by applying a threshold of 95% on the ANI, representing a 5% genetic distance cutoff. This threshold was chosen to group genomes with a high degree of sequence similarity while allowing for natural genomic variation within taxonomic groups;
3. <u>taxonomic label assignment</u>: each defined cluster was assigned with a taxonomic label based on a majority voting mechanism. This involved considering the NCBI taxonomic labels of all genomes within a cluster and selecting the most frequent label as the representative taxonomy for the entire cluster;
4. <u>handling taxonomic label duplicates</u>: in cases where two or more clusters share the same taxonomic label, we distinguish them as distinct clades assigning them an incremental numerical ID.

Among the set of clustered 26,285 reference genomes retrieved according to our extended definition of reference previously described, 686 genomes fell in clusters whose taxonomic label is different compared to the one reported in NCBI GenBank. Although MetaSBT does not utilize the taxonomic labels of MAGs as reported in NCBI GenBank, it is worth noting that among the set of 1,111 MAGs considered in the first update of the database, 1,051 genomes were misclassified in NCBI GenBank as reported in Supplementary Table S2.

It is also worth noting that the application of the widely adopted threshold of 5% on the genetic distance for the definition of species clusters during the generation of the first baseline of the MetaSBT database, led to the redefinition of 530 species each into a total of 3,236 clades, involving a total of 13,153 genomes. For instance, 2,456 genomes belonging to the *Rotavirus A* species have been reorganized into 217 different clades. The whole list of taxonomies affected by the *de novo* clustering of reference genomes is reported in the Supplementary Table S2.

It is also equally relevant that, subsequently to the clustering, 191 taxonomies disappeared because their genomes fell into clusters with a different taxonomic label assigned through the majority voting mechanism as previously mentioned above.

These findings are particularly relevant since there is a growing awareness on the importance of improving the classification of genomes in public databases, with different attempts to mitigate this problem proposed in the last few years [30,31].

### New clusters: real or artifacts?

The emergence of new clusters, populated by MAGs not assigned to any known species within the database, presents both exciting opportunities and interpretative challenges. While such clusters may represent actual novel taxa, it is essential to approach their interpretation with caution.

Several factors can contribute to the formation of new clusters that do not necessarily represent distinct species:

- <u>incomplete genome assembly</u>: MAGs, by their nature, are often incomplete representations of genomes. Clustering based on incomplete data can lead to artificial separation of closely related species, and the adoption of proper techniques for assessing the quality of such genomes is highly encouraged, as described early in this manuscript;
- <u>genome plasticity and horizontal gene transfer</u>: microbes exhibit varying degrees of genome plasticity, and horizontal gene transfer can confound phylogenetic analyses. Clusters may arise from the acquisition or loss of specific genomic regions rather than representing true species boundaries;
- <u>database coverage and bias</u>: the resolution of any taxonomic classification method is limited by the comprehensiveness and potential biases of the underlying database. Clusters composed of MAGs absent from reference databases may reflect gaps in our current knowledge rather than novel species.

Therefore, it is crucial to avoid prematurely assigning species-level classifications to new clusters solely based on their initial formation.

Longitudinal analysis, incorporating database updates and the addition of new MAGs, can provide valuable insights into the validity of new clusters:

- <u>cluster stability</u>: clusters that consistently expand with the addition of new data, demonstrating internal coherence and distinct boundaries from other clusters, are more likely to represent genuine taxonomic groups;
- <u>cluster quality</u>: a good quality of clusters in terms of completeness and contamination of their genomes, should also be taken into account while discriminating real clusters from potential artifacts;
- <u>comparative genomics</u>: in-depth comparative genomic analyses of MAGs within a cluster, focusing on core gene content, average nucleotide identity, and functional potential, can also provide further evidence against their classification as a novel species.

Looking back to Figure 2, among the 246 species clusters with a number of genomes >100 as reported in Figure 2, 207 are still unknown, with a number of MAGs ranging from 101 to 1,834 for the smallest and the biggest unknown species clusters respectively. Also note that 32 out of these 207 unknown species are high-quality clusters (i.e., in average, the completeness and contamination of the genomes in these clusters are >90% and ≤5% respectively), while 174 of them are medium-quality (i.e., completeness ≥50% and contamination ≤10%), and the remaining 1 is low-quality (i.e., completeness <50% and contamination ≤10%).

On the other hand, as we previously discussed, the vast majority of the species clusters defined by MetaSBT with a number of genomes ≤100 is still unknown. Among them, with a total of 32,353 clusters, 13,966 unknown species contain more than a single genome with 3,727 high-quality, 9,811 medium-quality, and 428 low-quality clusters according to the same thresholds of completeness and contamination defined above.

Other than excluding low-quality and single-sequence clusters, we cannot confirm, nor disprove, the authenticity of these clusters as real species yet. However, we should monitor their stability over future updates, as previously mentioned, to eventually confirm them as new candidate viral species.

Additionally, as reported in Supplementary Spreadsheet S2, it is worth to note that 46,649 MAGs (i.e., ∼19% of all processed MAGs) among those retrieved from NCBI GenBank and MGV, did not match with any of the clusters defined in MetaSBT (i.e., marked as *Unassigned*), not even at the kingdom level. Since the characterization of genomes in MetaSBT relies on clusters’ boundaries, and thus on the set of k-mers of all the genomes in the database, it does not necessarily mean that these genomes are not viruses. Note that the root node of the database tree contains the whole genetic content in terms of k-mers of all the genomes in the database. The set of k-mers of these unassigned and still uncharacterized genomes is not close enough to anything seen so far, and thus they must be reconsidered for future updates.

### Unlocking the quantitative profiling of unknown microbes in metagenomic samples with kraken2+bracken

Now that we have built a comprehensive database of viruses, we focus on benchmarking MetaSBT for the quantitative characterization of microbial populations starting from metagenomic samples.

Accurately characterizing the composition of microbial communities in metagenomic samples relies on our ability to identify and quantify the presence of both known and, crucially, unknown microbes. Traditional methods often struggle with the latter, but the integration of MetaSBT databases with *kraken2* [32] offers a powerful solution to rapidly detect unknown taxa in metagenomic samples.

*kraken* is a quantitative taxonomic classification tool that uses exact k-mer matching to rapidly and accurately assign taxonomic labels to metagenomic sequences. The following points summarize how it works:

1. <u>building a *kraken2* database</u>: a *kraken2* database is constructed from a set of genomes, representing known (and unknown in our case) microbial taxa. The database essentially stores all possible k-mers found within these genomes, along with their associated taxonomic information;
2. <u>classifying metagenomic reads</u>: when presented with a metagenomic sample, *kraken2* fragments the reads into their constituent k-mers. It then queries the database to identify which taxa are most likely to contain those specific k-mers;
3. <u>lowest common ancestor assignment</u>: *kraken2* employs a lowest common ancestor algorithm to assign taxonomic classifications. If a read’s k-mers match multiple taxa within the database, *kraken2* assigns it to the lowest taxonomic rank shared by all matches. This approach helps to account for uncertainties arising from shared genomic regions or incomplete database representation.

Using *kraken2* for benchmarking our databases comes with a series of advantages in terms of speed with extremely fast classifications, and flexibility of its databases that can be fully customized including new taxa as defined by MetaSBT.

Specifically, we used the taxonomic organization of all the 214,525 viral genomes in our MetaSBT database to build a custom *kraken2* database by running the *kraken* subroutine provided with MetaSBT which requires 1.2 GB of storage space only.

In order to benchmark the performance of our custom *kraken2* database, we focused on publicly available metagenomic samples from a study aimed to investigate the gut microbiome composition and metabolic activity in the context of Crohn’s disease (CD), ulcerative colitis (UC), and non-IBD (Inflammatory Bowel Diseases) controls across a US and Dutch cohort [33]. The dataset includes 220 paired-end fecal samples from the same number of distinct individuals, comprising 164 IBD patients—88 diagnosed with CD and 76 with UC—and 56 healthy controls.

*kraken2* was able to classify 4.32% of reads on average with our custom database compared to an average of 0.11% of reads classified with the classical *k2_viral* database (version dated 04/02/2025 available at https://benlangmead.github.io/aws-indexes/k2) built over known viral sequences only, as shown with the histogram in Figure 3C. Note that we applied a confidence threshold of 90% in *kraken2* for read classification and filtered input reads to retain only those with a minimum of PHRED quality score of 20. Within all the 220 metagenomic samples considered in this study, 219 viral species have been detected by *kraken2* in total, with 214 of them still unknown.

**Figure 3:**
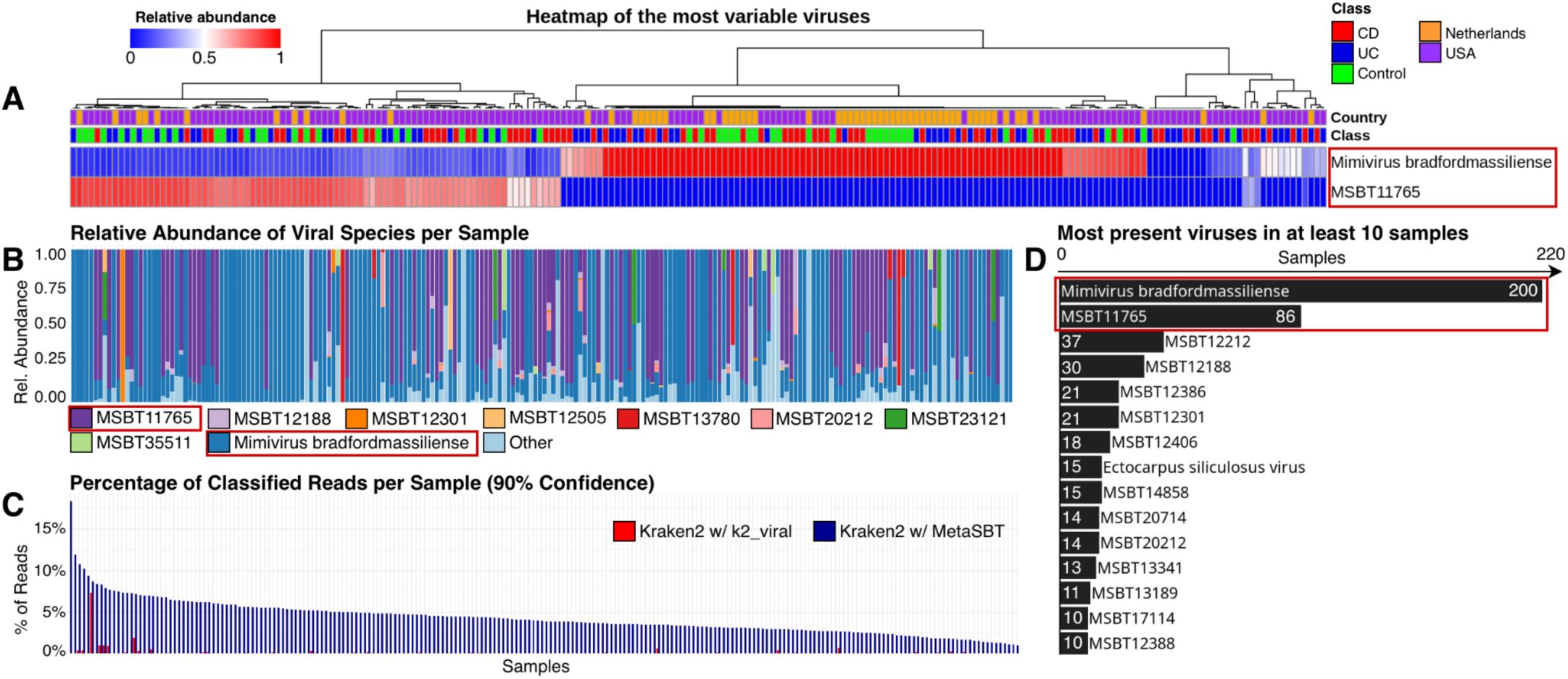
<u>Panel A</u> – heatmap of the most variable viruses in terms of their relative abundance estimated with *bracken*. Only the relative abundance of the first two most present species are reported. Samples are clustered according to the relative abundance of the viral species detected in the 220 samples; <u>Panel B</u> – percentage stacked bar chart showing the 9 most abundant viral species, plus an entry to groups all the remaining low abundant viruses (Other); <u>Panel C</u> – grouped bar chart with the percentage of classified reads per sample in descending order, highlighting the difference between the results produced by *kraken2* using the *k2_viral* database (red) and the MetaSBT enhanced database (blue); <u>Panel D</u> – histogram of the 10 most present viral species in the set of metagenomic samples considered in this study, showing the name of known and unknown viruses and the number of samples in which they have been detected by *kraken2*.

In order to check whether we could confidently state that the newly classified reads actually belong to viral species, we also performed the same analysis using the *k2_pluspfp* database (version dated 04/02/2025 available at the same link mentioned above) containing archaea, bacteria, viral, plasmid, protozoa, fungi, plant, and human sequences. Excluding the reads classified as viral, there are no overlaps between the reads classified using the two databases.

A list of the species detected in at least 10 samples are reported in Figure 3D. Note that the most prevalent species is known, i.e., *Mimivirus bradfordmassiliense*, and it appears in 200 out of 220 samples, followed by an unknown species, i.e., MSBT11765, which appears in 86 samples, which belongs to the unknown genus MSBT10778 and the known family *Graaviviridae*. This last one, according to ViralZone [34], is usually attributed to bacteriophages, having intestinal bacteria as their natural host.

It is also worth noting that only 8 of the 214 unknown viruses detected by *kraken2* are known at the genus level, with 3 of them belonging to the genus *Betapapillomavirus*, 3 → *Huchismacovirus*, 1 → *Coccolithovirus*, and 1 → *Primolicivirus*.

At the family level, 134 out of 214 unknown species are known, with 97 of them belonging to the family *Monoviviridae*, 28 → *Graaviviridae*, 3 → *Papillomaviridae*, 3 → *Smacoviridae*, 2 → *Phycodnaviridae*, and 1 → *Inoviridae*.

In terms of membership of unknown species to specific phyla, the majority of them belongs to the phylum *Pisuviricota* with 97 species, followed by *Uroviricota* → 39, *Negarnaviricota* → 27, *Taleaviricota* → 20, *Duplornaviricota* → 13, *Preplasmiviricota* → 4, *Cressdnaviricota* → 3, *Cossaviricota* → 3, *Hofneiviricota* → 3, *Nucleocytoviricota* → 3, *Dividoviricota* → 1, and *Peploviricota* → 1.

Based on our custom *kraken2* database, we also defined a *bracken* [35] database which can now easily estimate the abundance of known and unknown viral species in metagenomic samples as defined by MetaSBT. A percentage stacked bar chart showing the viral composition of the 220 samples according to the most abundant species detected by *bracken* is reported in Figure 3B. Note that the 2 most prevalent species highlighted in Figure 3D are also the most abundant ones.

Finally, we clustered the 220 samples based on the estimated abundance of all the 219 species detected by *kraken2* and quantified by *bracken* as shown with the heatmap in Figure 3A. Here we reported the two most variable viruses that better discriminate between two clearly distinct groups of samples, excluding the other 217 species that do not contribute to the clustering of samples at all because of their very low abundance. We also annotated the heatmap with information about the class of samples (i.e., CD, UC, and Control) in addition to the donors’ country (i.e., Netherlands and USA). Unfortunately, none of them contributes to explaining the clustering results, so there may be a hidden factor that has not been reported within the public samples metadata. Advancing hypotheses of superinfection exclusion or other infection mechanisms to explain our results would be purely speculative. These require additional analyses that fall out of the scope of our manuscript, while the goal of this section is to show how the integration of MetaSBT into common tools for microbiome analysis could lead to uncovering new insights from public studies.

### Galaxy: an ideal platform for MetaSBT

Galaxy [36] is the most popular web-based platform for reproducible and transparent bioinformatics analysis. It allows users to easily run software tools through a user-friendly graphical interface without the need to deal with command-line tools and complex software dependencies. Because of the amount of computational resources required to run MetaSBT on a massive number of genomes, our framework is not suitable for running on commodity hardware.

For these reasons, we integrated MetaSBT into the Galaxy platform to take advantage of the Galaxy computational infrastructure. It is available in the form of a tool suite comprising two different tools. The first one allows users to create a new database from scratch with the option to eventually update a predefined public MetaSBT database or a private user-defined one (see Figure 4 Panel A). On the other hand, the second tool allows users to profile genomes over a MetaSBT database at all the seven taxonomic levels, from the kingdom down to the species level, including the closest genome in the database (see Figure 4 Panel B). A list of software dependencies is reported in Table 4.

**Figure 4:**
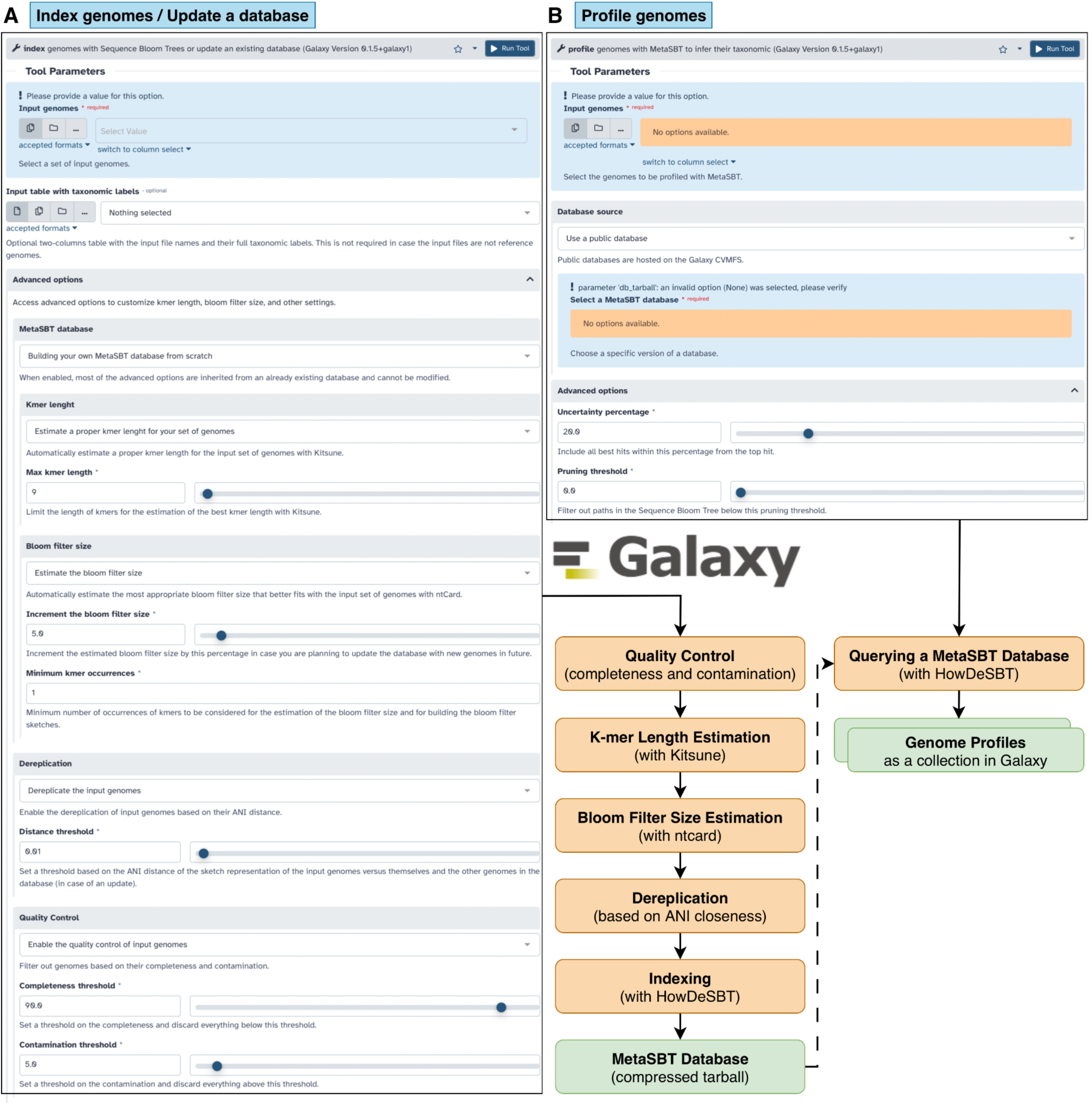
MetaSBT tool suite in Galaxy. <u>Panel A</u> shows the graphical user interface of the *index* tool that allows to index a set of genomes, and thus building a MetaSBT database, or update an existing MetaSBT database with new genomes, while <u>Panel B</u> shows the graphical user interface of the *profile* tool which allows to search for the closest taxonomic units and genomes in a given MetaSBT database.

**Table 4:** List of software dependencies and their version required to run MetaSBT in Galaxy.

| MetaSBT Tool | Process | Dependencies |
| --- | --- | --- |
| *index* | Quality Control<br>(<u>completeness</u> and <u>contamination</u>) | Kingdom dependent:<br>• Bacteria/Archaea → CheckM (≥1.2.4)<br>• Eukaryota → Busco (≥5.8.3)<br>• Viruses → CheckV (≥1.0.3) |
|  | K-mer Length Estimation | Kitsune (≥1.3.5) |
|  | Bloom Filter Size Estimation | ntcard (≥1.2.2) |
|  | Dereplication<br>(based on <u>ANI closeness</u>) | HowDeSBT (≥2.00.15) |
|  | Indexing |  |
| *profile* | Querying a MetaSBT Database | HowDeSBT (≥2.00.15) |

In order to allow users to incrementally update a public predefined MetaSBT database with their set of genomes, we stored our databases on the Galaxy distributed file system CVMFS [37]. Having our databases already in Galaxy is extremely useful so that users do not have to manually upload them into their working space, which could be problematic because of the large size of the databases. Please note that the MetaSBT public databases are also available outside of the Galaxy platform and maintained through the MetaSBT-DBs repository on GitHub at https://github.com/cumbof/MetaSBT-DBs/.

Furthermore, Galaxy’s collaborative features streamline the sharing of MetaSBT data. Researchers can easily grant collaborators access to private MetaSBT databases by simply sharing their Galaxy history, all without leaving the platform.

The integration of the MetaSBT tool suite into Galaxy workflows opens exciting possibilities for streamlining metagenomic analyses, like the workflows that already include the assembly of genomes from metagenomes. Its integration would allow the automatic incremental expansion of MetaSBT databases with newly assembled genomes, enabling the detection of novel species and strains. Such a capability would be particularly valuable in clinical settings or research focused on studying the microbial composition of samples related to specific diseases and health conditions, where identifying previously uncharacterized microbes could also lead to the definition of new biomarkers and eventually the development of personalized therapies.

## CONCLUSIONS

This work describes the first comprehensive database of viruses based on the MetaSBT framework. We classified many purported previously unknown viral species and demonstrated the importance of a careful curation of public databases to avoid misclassification of genomes. The new database and the application of MetaSBT for the quantitative characterization of microbiomes open the way to many new and exciting applications in the field of metagenomics.

We are fully committed to expanding the Viruses database by integrating additional publicly available datasets of genomes and sequences, and by eventually assembling new genomes from public metagenomes. We also plan to release new MetaSBT databases focused on other microbial kingdoms, including Bacteria, Archaea, and Fungi. Additionally, we are planning to enable the use of the Average Amino Acid Identity (AAI) in conjunction with the Average Nucleotide Identity (ANI) in future software versions. This will allow for more accurate clustering of genomes at higher taxonomic levels [38].

Building and maintaining these comprehensive databases presents significant challenges, particularly regarding the storage space required to keep track of all the bloom filters in the database tree that, in order to make the process of searching fast and efficient, is redundant by definition of sequence bloom tree. The integration of MetaSBT and its databases into the Galaxy platform helps address these challenges through the availability of scalable storage and computational resources for both building and updating the databases and for the profiling of genomes.

Moreover, the MetaSBT databases provide a powerful foundation for building custom databases for quantitative profilers like *kraken*. This opens exciting scenarios, such as reanalyzing large-scale metagenomic studies related to different diseases and health conditions that could lead to uncover previously undetected microbial biomarkers among still-unknown and yet-to-be-named species.

In conclusion, MetaSBT provides a powerful and scalable framework for characterizing microbial diversity, enabling the discovery and analysis of novel taxa, and ultimately advancing our understanding of the microbial world.

## ADDITIONAL INFORMATION

### Availability

MetaSBT’s code is open-source and available on GitHub at https://github.com/cumbof/MetaSBT (version 0.1.5) under the MIT license and installable as a Python 3.9 package from PyPI (*pip install metasbt*) and Bioconda [39] (*conda install -c bioconda metasbt*). Software documentation is available through the GitHub Wiki on the same repository. The MetaSBT Data Model is also defined in the same GitHub repository.

Public databases are hosted on the Galaxy CVMFS [37] and their metadata are also accessible through GitHub in the MetaSBT-DBs repository at https://github.com/cumbof/MetaSBT-DBs alongside the pipeline used to generate the databases in an incremental way. The custom *kraken* and *bracken* databases are also available in the same GitHub repository. Public databases are also indexed in Zenodo at https://doi.org/10.5281/zenodo.15580509.

Additionally, viral reference genomes and MAGs used for building the first two versions of the MetaSBT viral databases are available on the NCBI GenBank repository (Assembly Summary Report dated June 2024) at https://www.ncbi.nlm.nih.gov/genbank, while viral MAGs used for building the third incremental update of the MetaSBT viral database are available on the MGV repository (v1.0 dated July 8th, 2021) at https://portal.nersc.gov/MGV.

Gut microbiome samples here analyzed in the context of Crohn’s disease, ulcerative colitis, together with non-IBD (Inflammatory Bowel Diseases) samples, are publicly available on NCBI under the BioProject PRJNA400072.

Finally, the tool suite for the Galaxy Platform [36] is available on the official Galaxy ToolShed [40] after being reviewed and approved by the Intergalactic Utilities Commission, and it is fully integrated in the main instance of Galaxy https://usegalaxy.org/ and the Galaxy Microbiology Lab [41] instance https://microbiology.usegalaxy.eu/. A comprehensive tutorial about how to use the MetaSBT tool suite in Galaxy is also available on the Galaxy Training Network [42] at https://training.galaxyproject.org/training-material/topics/microbiome/tutorials/metasbt/tutorial.html.

### Author Contribution

FC and DB conceived the research; FC implemented the software, collected the reference genomes and metagenome-assembled genomes alongside sample metadata, and built the public databases; FC performed the analysis; DB supervised the research and provided critical feedback for the definition of the manuscript; FC and DB wrote the manuscript and both agreed with its final version.

### Conflict of Interests

DB has a significant financial interest in GalaxyWorks, a company that may have a commercial interest in the results of this research and technology. This potential conflict of interest has been reviewed and is managed by the Cleveland Clinic Foundation.

FC has no conflicts to disclose.

### Funding

This work has been supported by the National Institutes of Health [U24HG006620, U24CA231877].

## Acknowledgments

We would like to thank Robert S Harris and Paul Medvedev of the Department of Biochemistry and Molecular Biology of The Pennsylvania State University who developed HowDeSBT, the framework behind MetaSBT that is used for building Sequence Bloom Trees and indexing microbial genomes, for their support in expanding their software upon our requests. We would also like to thank the whole Galaxy Team, with special credits to Nate Coraor, for support in the integration of MetaSBT and its public databases into the Galaxy platform.

Finally, we would like to acknowledge the use of AI in refining the clarity and readability of this manuscript. The AI assistance was primarily used for tasks such as sentence restructuring, word choice suggestions, and identifying potentially unclear phrasing. We emphasize that the AI was used solely for language enhancement and did not contribute to the generation of research ideas, data analysis, or the interpretation of results. All conclusions drawn and insights presented in this manuscript are solely the product of the authors’ own analysis and expertise.

## Notes

https://github.com/cumbof/MetaSBT

https://github.com/cumbof/MetaSBT-DBs

### Competing Interest Statement

Daniel Blankenberg has a significant financial interest in GalaxyWorks, a company that may have a commercial interest in the results of this research and technology. This potential conflict of interest has been reviewed and is managed by the Cleveland Clinic Foundation.
Fabio Cumbo has no conflicts to disclose.

